# Domain-centric dissection and classification of prokaryotic poly(3-hydroxyalkanoate) synthases

**DOI:** 10.1101/693432

**Authors:** Zhanzhong Liu, Zuobin Zhu, Jianye Yang, Sheng Wu, Qinghua Liu, Mengmeng Wang, Huiling Cheng, Jiawei Yan, Liang Wang

**Author notes:** These authors contribute equally to the study (Zhanzhong Liu, Zuobin Zhu). Correspondence author: for all correspondence, please refer to Dr. Liang Wang.

## Abstract

Although many enzymes and multiple pathways involve in Polyhydroxyalkanoates (PHAs) synthesis, PHA synthases play a determinant role in the process, which include three subunits of PhaC, PhaE, and PhaR. Currently, PHA synthases are categorized into four classes according to its primary sequences, substrate specificity, and subunit composition. However, theoretical analysis of PHA synthases from the domain perspective has not been performed. In this study, we dissected PHA synthases thoroughly through analysis of domain organization. Both referenced bacterial and archaeal proteomes were then screened for the presence and absence of different PHA synthases along NCBI taxonomy ID-based phylogenetic tree. In addition, sequences annotated as bacterial and archaeal PhaCs in UniProt database were also analyzed for domain organizations and interactions. In sum, the *in-silico* study provided a better understanding of the domain features of PHA synthases in prokaryotes, which also assisted in the production of PHA polymers with optimized chemical properties.

## Introduction

Polyhydroxyalkanoates (PHAs) are a family of biodegradable and biocompatible polyesters that are synthesized intracellularly by a wide range of bacteria and archaea [1]. It naturally serves as carbon and energy resources for prokaryotes and provides important functions for their physiological activities, such as high salinity tolerance and desiccation resistance, *etc*., although the specific molecular mechanisms are still not solved. There are three types of PHAs according to their side chains, short-chain length (SCL) PHA (PHA_SCL_), medium-chain length (MCL) PHA (PHA_MCL_), and the mixture of both SCL and MCL PHAs (PHA_SCL-MCL_) [2]. Previous studies confirm that there are three enzymes involved in PHA synthesis, which include Acetyl-CoA acetyltransferase (PhaA), Acetoacetyl-CoA reductase (PhaB), and Poly(3-hydroxyalkanoate) polymerase subunit PhaC (PhaC) [3]. There are another two subunits, PhaE and PhaR, for Poly(3-hydroxyalkanoate) polymerase, which integrate with subunit PhaC to form active PHA synthase, respectively [4]. A heterogeneous group of small-sized proteins with no catalytic functions, Phasin (PhaP), was also tightly involved in granule structure formation and PHA metabolism [5]. As for PHA utilization, two enzymes, intracellular PhaZ and extracellular PhaZ, play important roles in this process [4]. Except for the above-mentioned classical pathway, enzymes from several other pathways are also indirectly involved in PHA synthesis, such as 3-oxoacyl-[acyl-carrier-protein] reductase (FabG), (R)-specific enoyl-CoA hydratase (PhaJ), Malonyl CoA-acyl carrier protein transacylase (FabD), Succinate-CoA ligase [ADP-forming] subunit alpha (SucD), NAD-dependent 4-hydroxybutyrate dehydrogenase (4HbD), and 4-hydroxybutyrate-CoA:CoA transferase (OrzF), *etc* [4]. Currently, more than 150 different hydroxyalkanoic acids are known to occur as constituents of PHAs [6] and a total of 14 PHA synthesis pathways have been identified with many enzymes involved in these processes [7].

The key enzymes involved in PHA synthesis are PHA synthases, which are a group of heterogeneous enzymes and include PhaC, PhaE, and PhaR that form homodimers, heterodimers, or work concordantly in their active states. These enzymes are currently divided into four categories, Class I, Class II, Class III and Class IV [8]. Class I, III and IV prefer SCL) carbon chain monomers (C3-5) while class II utilizes MCL PHA monomers (C6–14) [8]. In some exceptions, Class I PHA synthase could produce mixed PHAs incorporated with both SCL and MCL hydroxyalkanoates that show better functional properties compared with homo-polymers [2,7]. PhaC plays a key role in PHA synthesis and is the core subunit of PHA synthase. It has been extensively studied through metabolic engineering so as to optimize the properties of PHAs for industrial application. Active sites in different types of PhaCs belonging to a variety of bacteria were also reported [9]. However, these studies are only sporadic and cannot give an overview of PhaC features in prokaryotes. In addition, diversity and widespread of PhaC in bacterial species were only studied with a small set of genomes involved. PhaCs in archaea were recently investigated systematically. A novel type of Class III PhaC was widely identified with an elongated C terminus in halophilic archaea, which is indispensable for enzymatic activity [10]. Another study systematically investigated the distribution patterns of PHA synthesis pathway (PhaA, PhaB, and PhaC) and degradation pathway (PhaZ) in archaeal species, according to which PHA accumulation is skewedly associated with halophilic archaeal species [3].

Currently, the classification of PHA synthases is mainly through experimental studies based on primary sequences, substrate specificity, and subunit composition of the enzymes. In addition, PHA classes and corresponding descriptions recorded in the database are diverse with no consistent standards. One of the features of protein primary sequences used for identifying PhaCs is a lipase box of G-X-S-X-G [8]. In addition, a characteristic secondary structure containing α/β hydrolase fold was also highlighted [8]. However, these features are not sufficient to distinguish PhaCs in different classes. Due to the widespread accumulation of PHAs in prokaryotes, there is a large number of sequences of PHA synthases present in public database. However, there are no systematic studies looking at these sequences from domain-centric perspectives, which could provide a much clearer classification of and a better understanding about PHA synthases. For details of PHA synthases, please refer to Table 1.

**Table 1.**
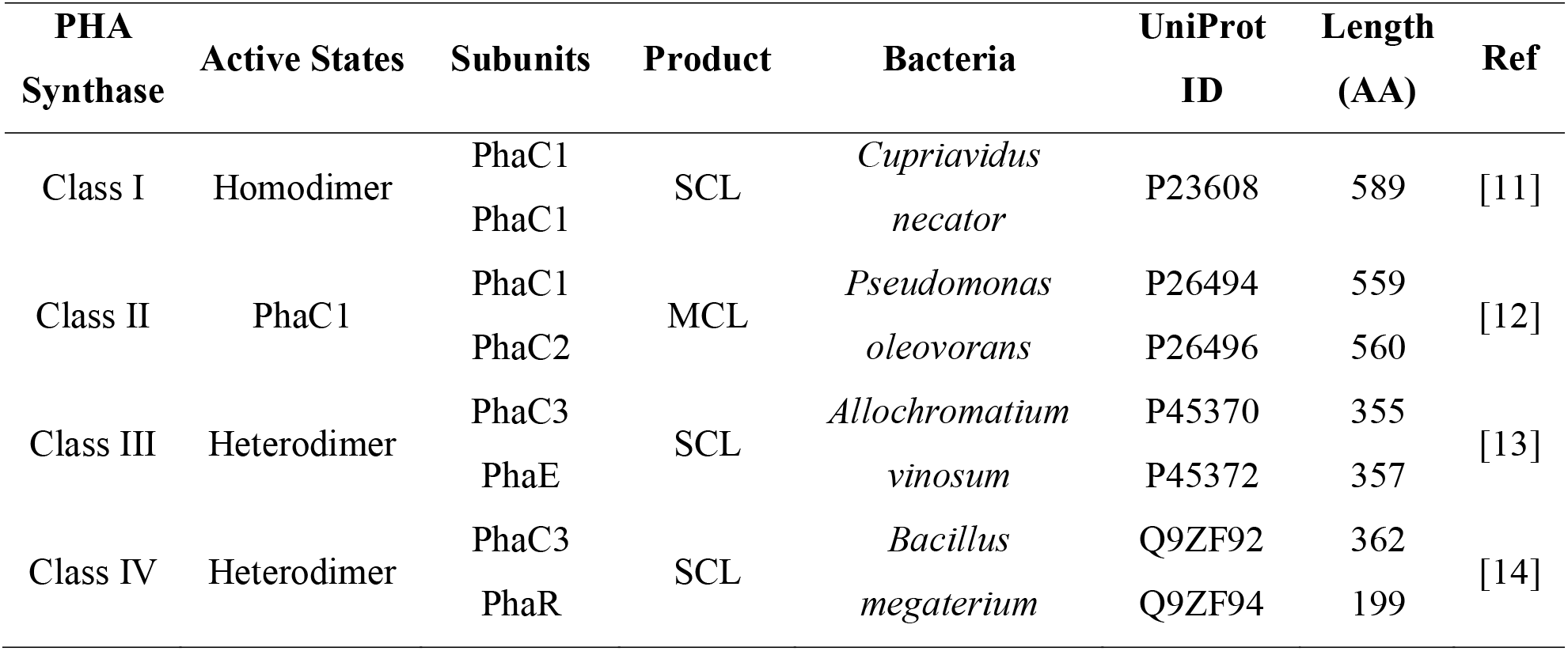
Summary of four classes of PHA synthases.

| PHA Synthase | Active States | Subunits | Product | Bacteria | UniProt ID | Length (AA) | Ref |
| --- | --- | --- | --- | --- | --- | --- | --- |
| Class I | Homodimer | PhaC1 | SCL | *Cupriavidus* | P23608 | 589 | [11] |
|  |  | PhaC1 |  | *necator* |  |  |  |
| Class II | PhaC1 | PhaC1 | MCL | *Pseudomonas* | P26494 | 559 | [12] |
|  |  | PhaC2 |  | *oleovorans* | P26496 | 560 |  |
| Class III | Heterodimer | PhaC3 | SCL | *Allochromatium* | P45370 | 355 | [13] |
|  |  | PhaE |  | *vinosum* | P45372 | 357 |  |
| Class IV | Heterodimer | PhaC3 | SCL | *Bacillus* | Q9ZF92 | 362 | [14] |
|  |  | PhaR |  | *megaterium* | Q9ZF94 | 199 |  |

From the domain-centric view, proteins can be considered as a combination of functionally and structurally independent domains [15]. Domains normally have independent evolutionary roadmaps, which can re-arrange and/or recombine with each other, leading to protein functional diversification and organism adaptation to new niches [15,16]. In addition, domain associations (co-occurrence) in a set of homologous sequences also reflect how proteins form and evolve at secondary structural level [15]. In this study, we first performed a systematic analysis of PHA synthases reported in literature via HMM models sourced from Pfam database (version 32.0, 17929 entries) and identified representative domains present in different classes. Their distributions in referenced prokaryotic proteomes were then studied in order to look for biologically meaningful patterns from evolutionary point of view. In addition, for sequences annotated as PhaCs that were retrieved from UniProt database, their domain interactions were also calculated and visualized to reflect its significance in different types of PhaCs.

In sum, this study looks into the diversity of PHA synthases from protein domain-based standpoint. This provides a novel angle for PHA synthase classification. In addition, we also overviewed the domain changes along taxonomy identifier based phylogenetic trees so as to understand how domain organization evolves to contribute to PHA synthesis. The conclusions would facilitate the understanding of the structures of PHA synthases and their evolutions in prokaryotes. It is also able to assist in the engineering efforts for the production of PHA polymers with optimized chemical properties.

## Methods and Materials

### Collections of sequences and proteomes in bacteria and archaea

Sequences of representative PHA synthases (PhaC, PhaE, and PhaR) were initially collected from literature (**Supplementary Table 1**), which were well classified into different groups based on corresponding phylogenetic studies [9,17–19] and were used as references to study domain organizations of PHA synthases in protein database [20]. Since UniProt Knowledgebase (UniProtKB) is the central-hub database for proteins with accurate, consistent and rich annotation, we choose it as our source for collecting high-quality data [21]. Two groups of PhaC-related sequences were selected from UniProt database that were present in bacteria and archaea, respectively. The sequences were picked-up via advanced searching of keywords “taxonomy:bacteria name:phac NOT name:phar NOT name:phae” and “taxonomy:archaea name:phac NOT name:phar NOT name:phae”. A total of 2004 bacterial proteins and 210 archaeal proteins were collected (**Supplementary Table 2**). As for referenced prokaryotic proteomes, 5447 bacterial species (**Supplementary Table 3**) and 287 archaeal species (**Supplementary Table 4**) were downloaded together with metadata from UniProt release 2019_02 [21].

**Table 2.** Summary of comparative distributions of PHA synthases in bacteria and archaea.

| PHA Synthase | Domain Organization | Pfam ID | Bacteria | Archaea |
| --- | --- | --- | --- | --- |
| PhaC | Abhydrolase_1 | PF00561 | + | + |
| PhaC | PhaC_N | PF07167 | + | - |
| PhaC | PhaC_N/Abhydrolase_1 | PF07167/ PF00561 | + | + |
| PhaC | Abhydrolase_1/HHH_5 | PF00561/PF14520 | - | + |
| PhaE | PHA_synth_III_E | PF09712 | + | + |
| PhaR | - | - | + | - |
| Novel PhaR | PHB_acc_N/PHB_acc | PF07879/PF05233 | + | - |
| PHB Polymerase | PHBC_N/PhaC_N | PF12551/PF07167 | + | - |

### Domain organizations and interactions in PHA synthases

PHA synthases from literature were initially dissected via HMMER webserver with default settings and the visualization of domain interactions were present in Figure 1 [22]. 17929 HMMs (version 32.0) were then downloaded from Pfam database for sorting domain organization of selected PhaCs from UniProt database [20,21]. Criterion for significant hits was set as E-value less than 1e-10 for all domains. Domain interactions in PhaCs selected from UniProt database were visualized by CytoScape based on Pfam-sourced HMMs, which gives the abundance of specific domains and also the associations among domains in a cluster of homologous sequences [23]. Abundance of a single domain is reflected by its circle radius while the strength of association between domains is displayed as line width. For domain circles, we artificially set the limits of circle sizes based on the least and the largest times of domain occurrence in collected sequences, while circle size of the rest domains are adaptively generated according to their ranks in the sort. In terms of domain associations, both the distances between domains and the times of domain co-occurrence were considered when calculating the connecting strength (Figure 2).

**Figure 1.**
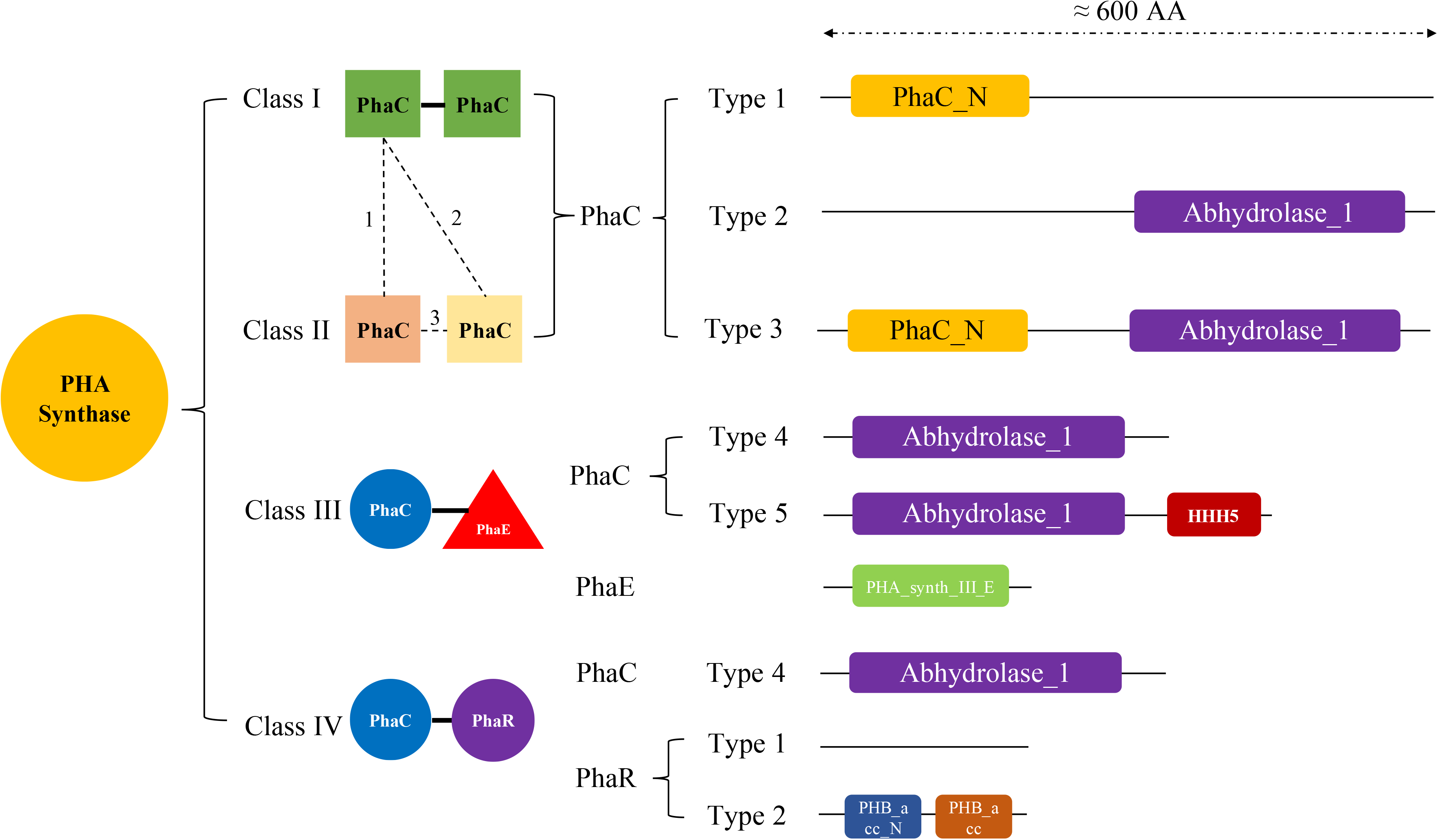
Illustration of four PHA synthase classification in prokaryotes. Class I PHA synthase is a PhaC homodimer while Class II PHA synthase is a heterodimer formed by two different PhaCs (*e.g.* combination 1, 2, 3 in dashed lines). Class III PHA synthase has two subunits PhaC and PhaE. Class IV PHA synthase consists of PhaC and PhaR. Based on domain organization and protein length, PhaC in four classes of PHA synthases is further divided into five sub-types as specified in the illustration.

**Figure 2.**
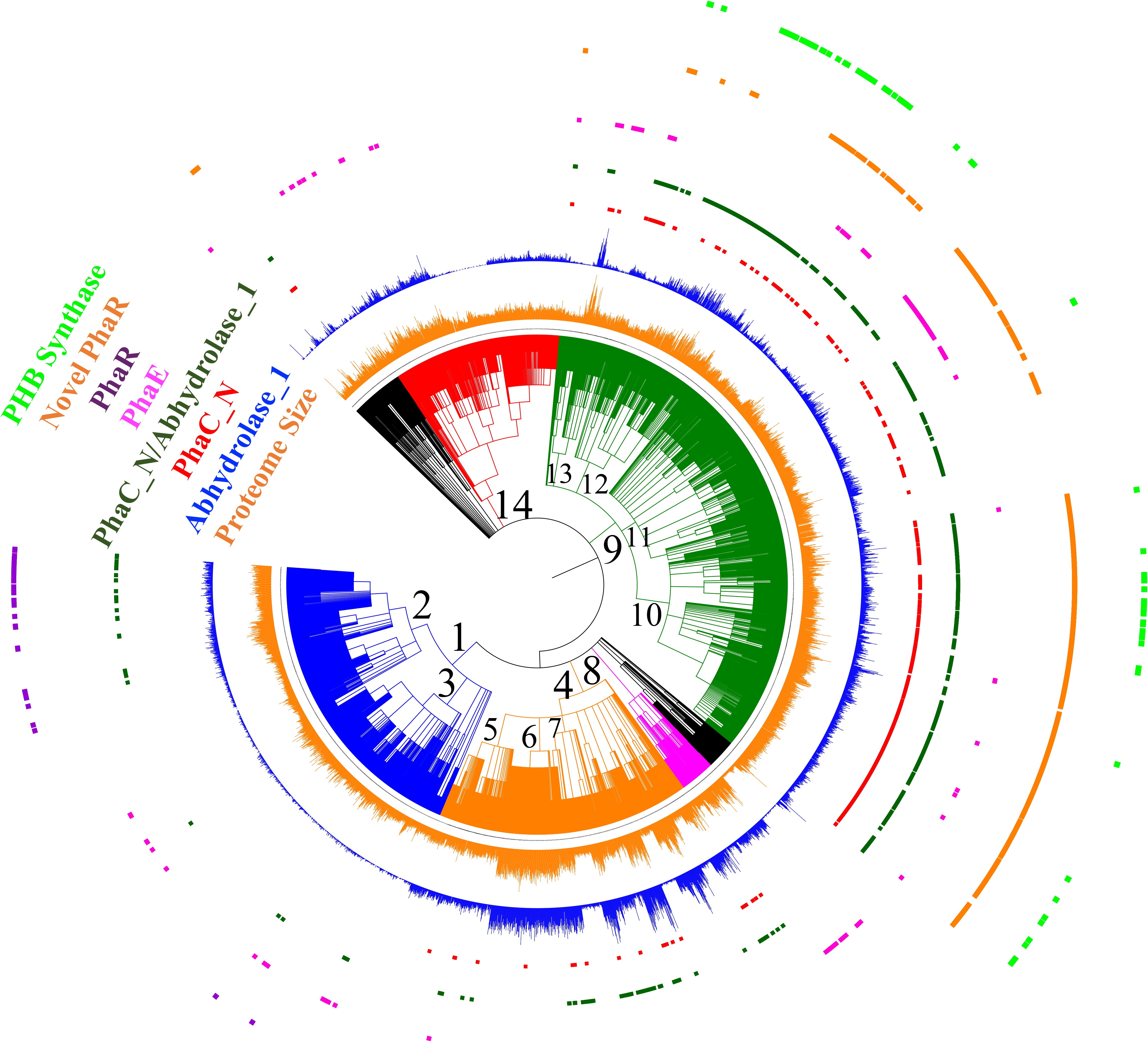
Distribution patterns of PHA synthases along NCBI taxonomy ID based phylogenetic trees. 5416 bacterial species sourced from UniProt database were included in the tree and belongs to 14 bacterial groups and classes, which are 1. *Firmicutes* (1239), 2. *Bacilli* (91061), 3. *Clostridia* (186801), 4. *Actinobacteria* (201174), 5. Micrococcales (85006), 6. *Streptomycetales* (85011), 7. *Corynebacteriales* (85007), 8. *Cyanobacteria/Melainabacteria* group (1798711), 9. *Proteobacteria* (1224), 10. *Alphaproteobacteria* (28211), 11. *Gammaproteobacteria* (1236), 12. *Betaproteobacteria* (28216), 13. Delta/Epsilon subdivisions (68525) and 14. FCB Group (1783270). A total of 7 groups of PHA synthases or subunits, together with bacterial proteome sizes were labelled and coloured accordingly in the phylogenetic tree.

### Homologs of PHA synthases in prokaryotic reference proteomes

According to preliminary analysis, PHA synthases (PhaC, PhaE, and PhaR) in literature exhibit a variety of domain organizations. Five subtypes of PhaC exist, which include type 1 with PhaC_N domain (PF07167), type 2 with Abhydrolase_1 domain (PF00561) in a long sequence, type 3 with PhaC_N and Abhydrolase_1 domains, type 4 with Abhydrolase_1 and HHH_5 domains (PF14520), type 5 with Abhydrolase_1 domain in a short sequence. As for subunit PhaE, it contains PHA_synth_III_E domain (PF09712). Subunit PhaR has two subtypes, type 1 with no domain and type 2 with PHB_acc_N domain (PF07879) and PHB_acc domain (PF05233). There is another group of PHA synthesis enzyme exclusively for PHB production that includes PHBC_N domain (PF12551) and PhaC_N domain, which is identified during HMM analysis. In order to identify homologs of PHA synthases in prokaryotic reference proteomes, 17929 HMMs were used to search selected bacterial and archaeal proteomes with E-value set to less than 1e-10. Due to the lack of known Pfam domain for some PhaR, local BLAST was performed by using PhaR (UniProt ID A0A386BX77) from *Bacillus cereus* as the query sequence with following criteria: E-value<1e-10, 30%<Similarity<100%, and 100 amino acids (AA) <subject length<400 AA. Results were then analyzed and mined through Python programming. Numbers of homolog hits, ranges of sequence lengths and E-values for each enzyme in each proteome were recorded in **Supplementary Table 3 and Supplementary Table 4**, respectively, which were then used for evolutionary analysis of PHA synthases by incorporating into phylogenetic trees.

### Phylogenetic analysis of PHA synthases

NCBI Taxonomy ID based phylogenetic trees, which were generated by phyloT https://phylot.biobyte.de/, were constructed to explore the distribution patterns of all different types of PHA synthases in collected bacterial and archaeal proteomes [24]. A maximum-likelihood (ML) phylogenetic tree based on a group of selected full length of PhaC sequences were constructed via fasttree, which were accompanied with domain organizations by in-house python scripts (available under request).

### Statistical analysis

All statistical analyses were performed by using R package. Student’s *t*-test was performed through two-tailed unequal variance with *P*-value less than 0.05 as significant difference.

## Results and discussion

### Domain-based classification of PHA synthases

It has been reported that class I PHA synthase is a homodimer of PhaC [8]. Class II PHA synthase is mainly found in *Pseudomonas* spp. and consists of two different PhaCs, PhaC1 and PhaC2, with PhaC1 as the active form [8]. In this study, based on the initial domain organizations of PHA synthases from literature, three sub-types of PhaCs in class I and II PHA synthases were identified and reported, that is, type 1 PhaC with only Abhydrolase_1 domain, type 2 PhaC with only PhaC_N domain, and type 3 PhaC with both PhaC_N and Abhydrolase_1 domains (**Supplementary Table 1**). According to the result, Class I PHA synthase has all three subtypes of PhaCs while Class 2 PHA synthase only harbours PhaC_N domain (type 2), which is not necessarily a general rule due to the small number of sequence samples. All three types of PhaCs have comparatively similar length of around 600 amino acids (AA). It would be interesting to find the unique features for the three sub-types of PhaCs. However, it is difficult to distinguish the two classes of PHA synthases just at the sequence level. Class III PHA synthase consists of PhaC and PhaE while class IV PHA synthase consists of PhaC and PhaR. PhaC in these two classes is shorter and around half length of PhaC in class I and II PHA synthases. Based on the domain analysis of the curated sequences from literature, PhaC in class III PHA synthases has two sub-types, one is with Abhydrolase_1 domain only and the other one is comprised of Abhydrolase_1 and HHH_5 domains. Interestingly, all the PhaCs with HHH_5 domain shared a high similarity with PhaCs with Abhydrolase_1 domain only and were found exclusively in halophilic archaea species such as *Haloarcula hispanica* and *Haloferax mediterranei, etc.* [18]. It has been proved that the extended C-terminus (HHH_5) of PhaC is essential for PHA production in halophilic archaea and could also play important roles in environment adaptation [10]. Please see Figure 1 for the illustrated classification of the four classes of PHA synthases.

A couple of other domains were also found in PHA synthases, which include DUF3141 and PHBC_N. DUF3141 resembles the Abhydrolase_1 domain and basically exists as a single domain protein that is mainly present in *Proteobacteria* [25,26]. It is also rarely associated with PhaC_N domain in several protein sequences according to Pfam database [20]. As for PHBC_N, it was also found to concatenate with PhaC_N to form a novel type of PhaCs, which does not fall into the four classes of PHA synthase and might have different functions in the PHB metabolism [26]. Unlike the sequence diversity of PhaC at the domain level, PhaE is comparatively conservative. All analyzed PhaE sequences consistently possess the PHA_synth_III_E domain. In contrast, there is no domain information for PhaR sequences sourced from *Bacillus* related species, such as *Bacillus cereus* and *Bacillus megaterium*. However, a novel type of PhaR from other bacteria such as *Chromohalobacter salexigens* and *Bradyrhizobium japonicum* consists of PHB_acc_N (PHB/PHA accumulation regulator DNA-binding domain) and PHB_acc (PHB accumulation regulatory domain), which suggested that PhaR sequences could also be rather diverse and different types of PhaR might arise multiple times in evolution with similar functions.

### Distribution patterns of PHA synthases

Due to the diversification of PHA synthases and its subunits, understanding how these enzymes distribute along phylogenetic trees is necessary to better interpret their functions, domain organizations and evolution. According to the domain-based classification of PHA synthases, we systematically searched 5447 bacterial species and 287 archaeal species for how PHA synthases distribute along phylogenetic trees, respectively.

#### Bacteria

In total, only 5416 bacterial species were mapped in the phylogenetic tree due to the outdated taxonomy IDs, which were mainly divided into 14 bacterial groups and classes. These include (1) *Firmicutes*, (2) *Bacilli*, (3) *Clostridia*, (4) *Actinobacteria*, (5) *Micrococcales*, (6) *Streptomycetales*, (7) *Corynebacteriales*, (8) *Cyanobacteria/Melainabacteria* group, (9) *Proteobacteria*, (10) *Alphaproteobacteria*, (11) *Gammaproteobacteria*, (12) *Betaproteobacteria*, (13) Delta/Epsilon subdivisions and (14) the FCB group (Figure 2). Based on the dissection of HMM domain models for all the sequences in those proteomes, a total of 7 groups of PHA synthases and/or subunits, were present. PhaC has three subclasses, sequences with only Abhydrolase_1 domain, only PhaC_N domain or both Abhydrolase_1 and PhaC_N domains. According to the result, the most abundant subtype of PhaC should be those with Abhydrolase_1 domain. Abhydrolase_1 belongs to the superfamily of hydrolytic enzymes that share a common fold, which has been identified in a large collection of enzymes with different origins and functions [27]. Thus, Abhydrolase_1 is rather abundant in bacterial proteomes. Sequences with only Abhydrolase_1 domain have two sub-types, long and short. Long PhaC with only Abhydrolase_1 domain should belong to Class I and Class II PHA synthases while short PhaCs with only Abhydrolase_1 domain belong to Class III and Class IV PHA synthases. However, due to the difficulty to determine the boundaries between long and short PhaCs, it is hard to show how the two subtypes distribute along phylogenetic trees. In contrast, Class III and Class IV PHA synthases are determined by the presence of PhaE and PhaR. For the other two types of PhaCs with domains of PhaC_N or PhaC_N/Abhydrolase_1, they are mainly clustered in the phylum of *Actinobacteria* and *Proteobacteria* while species in other groups or classes only sporadically harbour the enzymes. Thus, the two types of PhaCs might serve special functions in these two phylums and requires further experimental investigation for their roles

PhaE is an independent subunit of Class III PHA synthases. According to our study, PhaE is sporadically distributed along the phylogenetic tree, showing no preferred association with bacterial species in any phylums or classes. However, it is noteworthy that a protein (UniProt ID A0A252E8H1) with both Abhydrolase_1 and PHA_synth_III_E domains was found in *Nostoc* sp. T09, a genus of *Cyanobacteria* that has been found in various environments. This novel type of enzyme has never been reported before and could be caused by a gene fusion event in the bacteria or could also be a sequencing error during genome assembly. Further investigation is required to address this issue. If indeed a novel protein exists that harbors functions of both PhaC and PhaE, it would be interesting to see what the properties and kinetics of this enzyme are for PHA synthesis when compared with the PhaC and PhaE heterodimer.

PhaR is the subunit of Class IV PHA synthases. It was found to be specifically associated with *Bacilli* class in the *Firmicutes* Group, as previous studies suggested [14]. According to the domain analysis, another two species, *Desulfallas gibsoniae* DSM 7213 (E-value=1.86E-17) and *Thermosyntropha lipolytica* DSM 11003 (E-value=2.39e-14), also possess PhaR. Thus, this enzyme might play important roles in these bacterial species. By figuring out how and why they acquire PhaR genes, it could be helpful for us to better understand PHA metabolism in bacteria. Interestingly, a novel group of PhaR with two small domains, PHB_acc_N and PHB_acc, were found to be widely and abundantly distributed in the subdivisions of *Proteobacteria* class. In addition, a new PHB synthase, which has both PHBC_N and PhaC_N domains, were mainly found in alpha- and beta-*Proteobacteria* subdivisions. Their physiological functions and distribution patterns are worthy of further exploration so as to better understand their unique roles.

#### Archaea

Previous bioinformatics analysis showed the complete metabolism pathway of PHA in archaea by focusing on enzymes including PhaA, PhaB, PhaC, PhaE, PhaP, and PhaZ, according to which it was confirmed that PHA accumulation is tightly related to halophilic archaea with larger proteome size and higher GC contents [3]. In this study, we are only interested in PHA synthases (PhaCs and its subunits) and explored their distribution patterns in archaea. It was shown that PhaCs with single Abhydrolase_1 domain are most abundant in archaeal species, similar to the pattern in bacteria, although some of the strains do not harbour the enzyme, such as *Nanoarchaeum equitans* and *Ignicoccus hospitalis*, *etc*. Interestingly, *Ignicoccus hospitalis* is the host organism for *Nanoarchaeum equitans* [28]. In addition, species in unclassified Euryarchaeota such as candidate divison MSBL1 archaeon seems to have very small genomes, most of which tend to lack PhaCs, losing PhaC accumulation ability. Since the uncultured archaeal lineage MSBL1 strains are from brine pool, they are also adapted to hypersaline environment [29]. Thus, PHA accumulation might not be necessary for saline environment adaptation and survival of halophilic archaea. On the other hand, *Halobacteria* shows to possess multiple copies of PhaCs with only a single Abhydrolase_1 domain. As for the second type of PhaCs that consists of two domains, PhaC_N and Abhydrolase_1, they are mainly spread in the *Halobacteria* phylum and TACK group. Interestingly, there is no PhaC in archaea that has a single domain of PhaC_N. Thus, PhaC with only PhaC_N domain could be a marker for PHA synthases in bacteria and might play unique physiological features in PHA synthesis. A novel type of PhaC that is solely identified in archaea have two domains, Abhydrolase_1 and HHH_5, as illustrated above (Figure 1). The enzyme is mainly restricted to *Halobacteria* except for one strain uncultured archaeon A07HB70. Class III PHA synthase is mainly found in *Halobacteria* and the TACK group due to the presence of PhaE while Class IV PHA synthase is not found in archaea, which reflects the unique evolutionary features of archaeal species when compared with bacterial species. It is worth noting that both *Thaumarchaeota* phylum from the TACK group and the Halobacteria class seem to have abundant and diverse PHA synthesis enzymes.

In order to better understand the distribution patterns of PHA synthases in both bacteria and archaea, we summarized the corresponding presence and loss of PHA synthase types in Table 2. For each domain, its Pfam ID is also shown so its functions could be easily retrieved from Pfam database.

### Domain interactions (co-occurrence) in PhaCs

Distribution patterns of PHA synthases give us a clear view of how different types of enzymes evolve and cluster along phylogenetic trees. However, it cannot measure domain abundance and co-occurrence quantitatively in a group of protein with multiple classes, which could reflect core domains and domain relationships in a clustered sequence group [15]. In this study, PhaC sequences were collected from UniProt database as long as their protein names include PhaC and does not include PhaE and PhaR. Through a thorough analysis of all selected sequences, we visualized the domain interactions in both archaeal PhaCs (Figure 4) and bacterial PhaCs (Figure 5).

**Figure 3.**
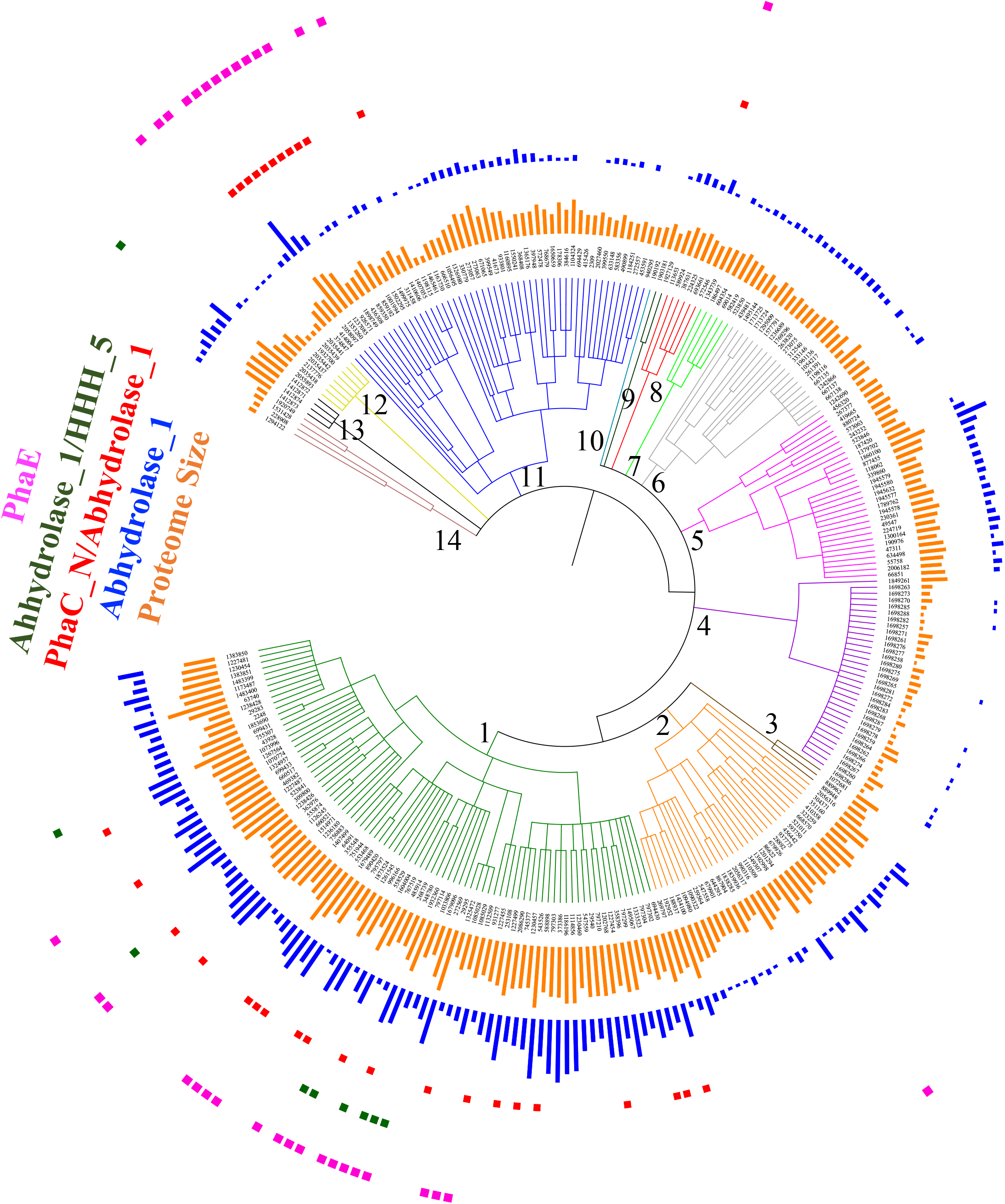
Distribution patterns of PHA synthases along NCBI taxonomy ID based phylogenetic trees in 287 archaeal species sourced from UniProt database. A total of 14 archaeal groups and classes were annotated, which include 1. *Halobacteria* (183963), 2. *Methanomicrobia* (224756), 3. Candidatus *Nanohaloarchaeota* (1462430), 4. Unclassified *Euryarchaeota* (33867), 5. *Methanomada* Group (2283794), 6. *Diaforarchaea* Group (2283796), 7. *Thermococci* (183968), 8. *Archaeoglobi* (183980), 9. *Methanonatronarchaeia* (171536), 10. *Methanopyri* (183988), 11. TACK group (1783275), 12. Unclassified Archaea (29294), 13. Environmental Samples (48510), and 14. DPANN Group (1783276). PHA synthases or subunits, together with archaeal proteome sizes, were shown in the phylogenetic tree and were annotated accordingly.

**Figure 4.**
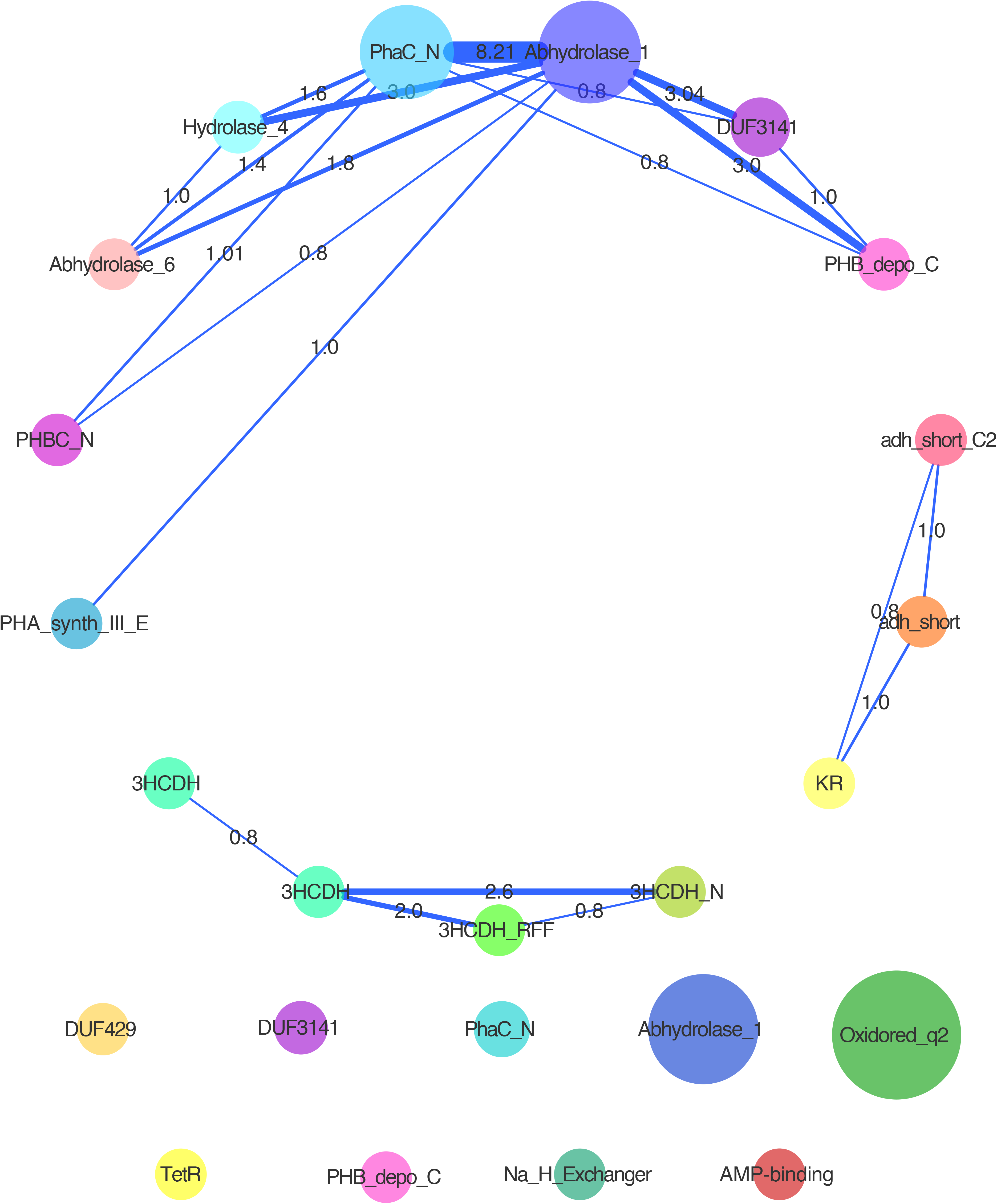
Interactions of domains in 2004 annotated PhaC enzymes from bacteria. All sequences were thoroughly collected from UniProt database. Radius of color-filled circles reflects the comparative abundance of the domain in all analyzed sequences. Edge shows the connection of domain pairs and its width indicates strength of the interactions between the two domains.

**Figure 5.**
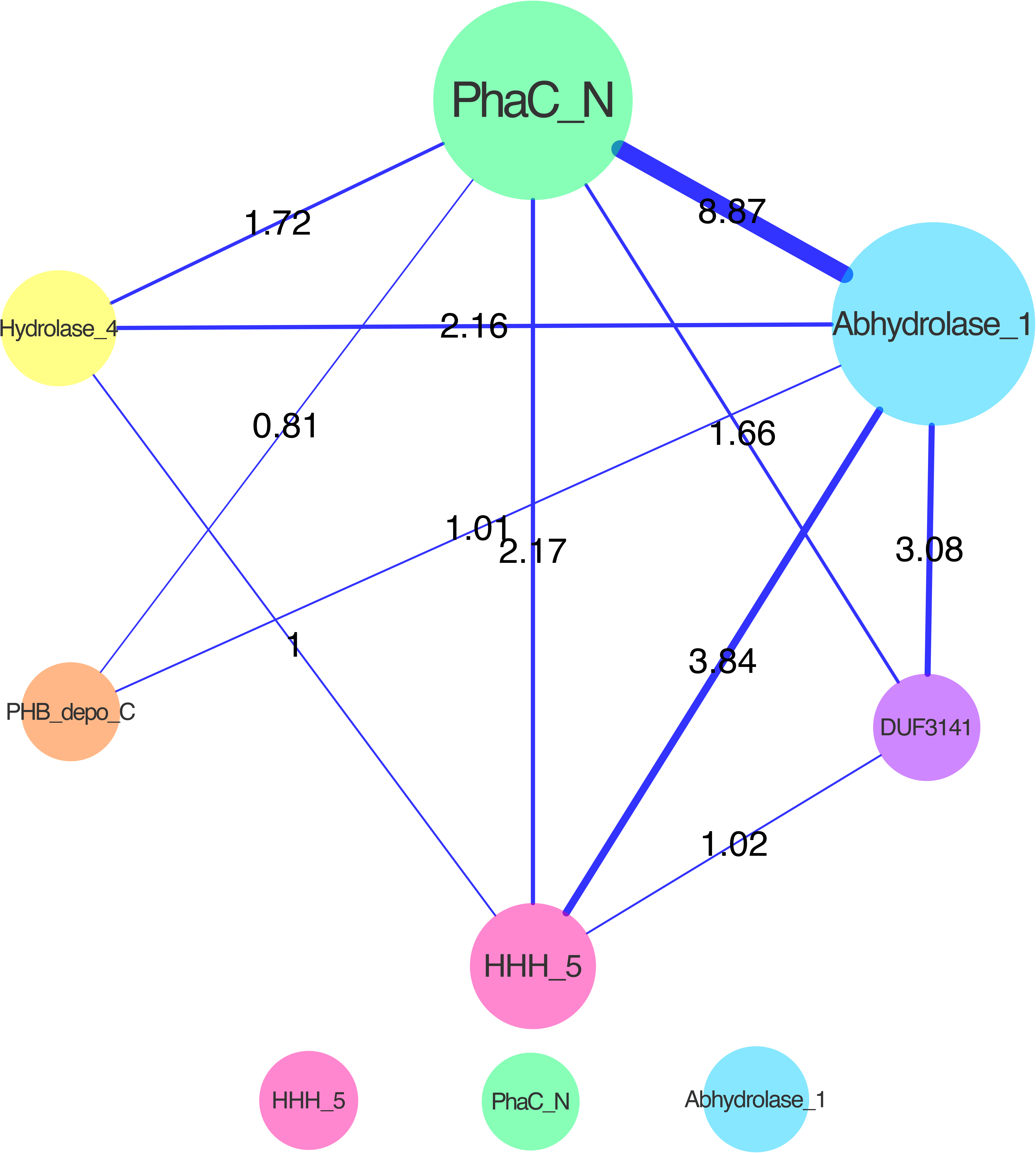
Interactions of domains in 210 annotated PhaC enzymes from archaea. Six domains, PhaC_N, Abhydrolase_1, HHH_5, Hydrolase_4, PHB_depo_C and DUF3141, were present. Domain abundance and connection strength were illustrated through circle radius and line width.

As for bacterial PhaCs, it is obvious that all domains generated through HMM dissection were divided into four groups, among which three groups are independently clustered. The fourth group include single domains with no co-occurrence with others, though they could also be present in the clustered groups and interact with other domains (Figure 4A). The dominant group includes domains of Abhydrolase_1, PhaC_N, DUF3141, PHBC_N, Hydrolase_4, PHB_depo_C, Abhydrolase_6 and PHA_synth_III_E, among which Hydrolase_4, PHB_depo_C, and Abhydrolase_6 were not identified in the four PHA synthase classes. PHB_depo_C is the C-terminal domain of bacterial poly(3-hydroxybutyrate) depolymerase that degrades PHB into oligomers and monomers of 3-hydroxy-butyric acid [30]. Since PHB_depo_C is dominantly present in intracellular PHB depolymerase, it is interesting to see that it also co-occurs with Abhydrolase_1 in sequences annotated as Poly-beta-hydroxybutyrate polymerase or Polyhydroxyalkanoate synthase. In addition, association between PHB_depo_C and DUF3141 (resembled to Abhydrolase_1) was observed only in a protein named DUF3141 domain-containing protein (UniProt ID A0A3A4PYR6) in the strictly anaerobic *Desulfobacteraceae bacterium*, which was further confirmed through whole UniProt database search. As for PHB_depo_C (E-value= 9.8e-13), Abhydrolase_1 (E-value=3.5e-16), and PhaC_N (E-value=8.8e-11), their associations were identified in a protein titled Class III poly(R)-hydroxyalkanoic acid synthase subunit PhaC (UniProt ID A0A1Q7IEF1) in *Ktedonobacter* sp. These associations deserve further investigation both theoretically and experimentally in order to understand their origin and potential functions.

As for the other two clustered domains and some of the independent domains such as Oxidored_q2 and Na_H_Exchanger, their existence in the domain interaction map largely indicates mis-annotations that might be present in UniProt database. For example, the adh_short, adh_short_2, and KR cluster is mainly found in 3-oxoacyl-ACP reductase that catalyses the 3-oxoacyl-ACP reduction step in the fatty acid synthesis pathway [31]. However, in *Burkholderia puraquae* and *Burkholderia pyrrocinia*, proteins (UniProt ID A0A1D8BIL9 and A0A0N9MGN9) with adh_short domain were named Acetoacetyl-CoA reductase and PhaC, respectively. As for 3CDH related domain cluster, it is mainly present in 3-hydroxyadipyl-CoA dehydrogenase. However, enzymes with the 3CDH domains in *Desmospora* sp. (UniProt ID F5SKZ9), *Pseudomonas entomophila* (UniProt ID Q1I9V2), *Enterobacter hormaechei* (UniProt ID F5RRQ1) were all annotated as 3-hydroxyacyl-CoA dehydrogenase PhaC.

In terms of archaeal PhaCs, their domain organizations showed that a total of six domains, PhaC_N, Abhydrolase_1, HHH_5, Hydrolase_4, PHB_depo_C and DUF3141, were present and all connected together (Figure 5). In addition, PhaC_N and Abhydrolase_1 shows the strongest connections. Except for Hydrolase_4, all domains have been observed in PHA synthase and all the domains co-occurs with each other to form a single cluster, though HHH_5, PhaC_N, and Abhydrolase_1 are also able to exist independently in single domain form. Proteins (UniProt ID A0A062VD38 and A0A062V164) from archaea Candidatus *Methanoperedens nitroreducens* harbour PHB_depo_C, Abhydrolase_1 and PhaC_N, which also raised the question of why domains for synthesis and degradation co-occur. In addition, it suggested that this type of enzyme is not restricted to bacteria and should be explored further, especially for the functions that the PHB_depo_C domain might play in the enzyme.

## Conclusion

PHAs are a group of biodegradable and biocompatible natural thermoplastics with good mechanical properties. A variety of building blocks are available for PHA synthesis, which leads to its structural polydispersity. PHA synthases play a central role in PHA biosynthesis. Its activity and substrate specificity determine characteristics of the generated polymers. Although PHA synthases are currently divided into four classes based on kinetics and mechanisms of reactions, no studies attempt to classify and summarize PHA synthases via domain-centric viewpoints. In this study, we re-defined five sub-types of PhaCs in the four classes of PHA synthases according to domain organizations, together with the conservative PhaE and the two types of PhaR. Another two rare types of PhaCs, one with PHA_synth_III_E domain and the other one with PHB_depo_C domain, were also observed and required further investigation for their existence and functions. In addition, distributions of different types of PHA synthases along evolutionary trees were also explored. Interesting patterns were identified with insights into how PHA synthases were preferred in different bacterial and archaeal species, which seem to have different preferences for PHA synthases and might be determined by unique evolutionary pathways and adaptation to specific niches. In addition, we also investigated the domain co-occurrence in PhaC sequences collected from UniProt database. Core domains and correlations among domains were studied. However, misannotation of PhaC enzymes from UniProt database should draw some attention and information from public database should be used with carefulness. In sum, through domain-based analysis, we get a better understanding and an overview of how PHA synthase is classified, which also give hints to further industrial engineering of the enzymes to develop better bioplastic products.

## Supplementary Materials

Table S1: Domain organizations of phylogenetically classified enzymes directly associated with PHA synthesis that include PhaC, PhaE, and PhaR; Table S2: Collection of bacterial and archaeal protein sequences annotated as PhaCs in UniProt database; Table S3: Distributions of PHA synthase related domains in 5447 bacterial proteomes; Table S4: Distributions of PHA synthase related domains in 287 archaeal proteomes.

## Author Contributions

Conceptualization, Liang Wang; Data curation, Qinghua Liu and Mengmeng Wang; Formal analysis, Zhanzhong Liu, Zuobin Zhu and Liang Wang; Funding acquisition, Liang Wang; Methodology, Liang Wang; Project administration, Liang Wang; Validation, Jianye Yang, Sheng Wu and Huiling Cheng; Visualization, Zhanzhong Liu and Zuobin Zhu; Writing – original draft, Liang Wang; Writing – review & editing, Zhanzhong Liu, Zuobin Zhu and Jiawei Yan.

