## Supplementary Table S1 for "Domain-centric dissection and classification of prokaryotic poly(3-hydroxyalkanoate) synthases"

**Supplementary Table 1** Domain organizations of phylogenetically classified enzymes directly associated with PHA synthesis that include PhaC, PhaE, and PhaR.

| **Species** | **UniProt ID** | **PHA Synthase** | **PhaC Class** | **Pfam Domain** | **Ref** |
| --- | --- | --- | --- | --- | --- |
| *Acinetobacter* sp. | Q57164 | PHA synthase | I | PhaC_N | [1] |
| *Aeromonas hydrophila* | Q8L1Y0 | PHA synthase | I | PhaC_N | [1] |
| *Aeromonas punctata* | O32471 | PHA Synthase | I | PhaC_N | [1] |
| *Agrobacterium tumefaciens* CFBP 5771 | A0A1S7SU53 | Poly-beta-hydroxybutyrate polymerase | I | PhaC_N  Abhydrolase_1 | [1] |
| *Alcaligenes* sp. | P94189 | PHA synthase | I | PhaC_N | [1] |
| *Allochromatium vinosum* DSM 180^*^ | P45370 | Poly(3-hydroxyalkanoate) polymerase subunit PhaC | I | Abhydrolase_1 | [1] |
| *Aromatoleum aromaticum* EbN1 | Q5NZ38 | PHA synthase subunit PhaC | I | Abhydrolase_1 | [2] |
| *Aromatoleum aromaticum* EbN1 | Q5P1L8 | Poly-beta-hydroxybutyrate polymerase | I | PhaC_N | [2] |
| *Aromatoleum aromaticum* EbN1 | Q5P3D9 | Putative poly-beta-hydroxybutyrate synthase | I | PhaC_N  Abhydrolase_1 | [2] |
| *Azohydromonas lata* | Q9Z3D5 | Poly-beta-hydroxybutyric acid synthase | I | PhaC_N | [1] |
| *Azohydromonas lata* | Q9ZB54 | Polyhydroxyalkanoate synthase | I | PhaC_N | [3] |
| *Azorhizobium caulinodans* DSM 5975 | O66392 | Poly(3-hydroxyalkanoate) polymerase subunit PhaC | I | PhaC_N  Abhydrolase_1 | [1] |
| *Bradyrhizobium diazoefficiens* IAM 13628 | Q89NV2 | Poly-beta-hydroxybutyrate polymerase | I | PhaC_N | [4] |
| *Bradyrhizobium japonicum* | A0A1Y2JQJ1 | Class I poly(R)-hydroxyalkanoic acid synthase | I | PhaC_N | [4] |
| *Burkholderia cepacia* | G3M4I8 | Polyhydroxyalkanoate synthase | I | PhaC_N  Abhydrolase_1 | [1] |
| *Burkholderia cepacia* | A0A2X2S146 | Class I poly(R)-hydroxyalkanoic acid synthase | I | PhaC_N | [4] |
| *Burkholderia* sp. DSM 9242 | Q9RB82 | PhaC | I | PhaC_N | [1] |
| Caulobacter vibrioides (Caulobacter crescentus) | Q9F4K5 | PHB synthase | I | PhaC_N | [1] |
| *Chromobacterium violaceum* DSM 30191 | Q9ZHI2 | Poly(3-hydroxyalkanoate) polymerase | I | PhaC_N | [1] |
| *Crenarchaeota archaeon* | A0A370LKH3 | Class III poly(R)-hydroxyalkanoic acid synthase subunit PhaC | I | Abhydrolase_1 | [1] |
| *Cupriavidus necator* DSM 428 | Q0KA68 | Poly(3-hydroxybutyrate) polymerase | I | PHBC_N PhaC_N | [3] |
| *Cupriavidus necator* DSM 428 | P23608 | Poly(3-hydroxyalkanoate) polymerase subunit PhaC | I | PhaC_N | [1] |
| *Delftia acidovorans* | O87110 | PHA synthase | I | PhaC_N | [1] |
| *Halomonas* sp. O-1 | A0A060NBA3 | Polyhydroxyalkanoate synthase | I | PhaC_N  Abhydrolase_1 | [2] |
| *Halomonas elongata* DSM 2581 | E1V6A8 | Poly-beta-hydroxyalkanoate polymerase | I | PhaC_N | [2] |
| *Halomonas* sp. TD01 | F7SP57 | Putative poly(3-hydroxyalkanoate) synthetase | I | DUF3141 | [2] |
| *Halorhodospira halophila* SL1 | A1WYS5 | Poly(R)-hydroxyalkanoic acid synthase, class I | I | PhaC_N | [2] |
| *Methylobacterium extorquens* | P52070 | Poly(3-hydroxyalkanoate) polymerase | I | PhaC_N  Abhydrolase_1 | [1] |
| *Novosphingobium aromaticivorans* DSM 12444 | Q2GB25 | Poly(R)-hydroxyalkanoic acid synthase, class I | I | PhaC_N  Abhydrolase_1 | [4] |
| *Paracoccus denitrificans* | Q9WX80 | Poly (3-hydroxyalkanoate) synthase | I | PhaC_N  Abhydrolase_1 | [1] |
| *Pseudomonas* sp. 61-3 | Q9Z3Y3 | PHB synthase | I | PhaC_N | [1] |
| *Rhizobium etli* CFN 42 | Q52728 | Poly(3-hydroxyalkanoate) polymerase subunit PhaC | I | PhaC_N | [1] |
| *Rhodobacter capsulatus* NBRC 16581 | D5APD9 | Poly(3-hydroxyalkanoate) polymerase | I | PhaC_N | [1] |
| *Rhodobacter sphaeroides* sp. denitrificans | A0A370Y039 | Class I poly(R)-hydroxyalkanoic acid synthase | I | PhaC_N | [1] |
| *Rhodospirillum rubrum* | Q9RNU7 | Polyhydroxyalkanoate synthase | I | PhaC_N | [1] |
| *Rickettsia prowazekii* Madrid E | Q9ZCD7 | Poly-beta-hydroxybutyrate polymerase ( PhbC2） | I | PhaC_N | [1] |
| *Rickettsia prowazekii* Madrid E | Q9ZCJ4 | Poly-beta-hydroxyalkanoate polymerase ( PhbC1） | I | Abhydrolase_1 | [1] |
| *Rhizobium meliloti* 1021 (*Sinorhizobium meliloti*) | P50176 | Poly(3-hydroxyalkanoate) polymerase subunit PhaC | I | PhaC_N | [1] |
| *Synechocystis* sp. PCC 6803^*^ | P73390 | Poly(3-hydroxyalkanoate) polymerase subunit PhaC | I | Abhydrolase_1 | [1] |
| *Thiococcus pfennigii* (*Thiocapsa pfennigii*) | A0A060Q549 | PHA synthase | I | Abhydrolase_1 | [1] |
| *Thiocystis violacea*^*^ | P45366 | Poly(3-hydroxyalkanoate) polymerase subunit PhaC | I | Abhydrolase_1 | [1] |
| *Vibrio parahaemolyticus* | Q938U7 | Polyhydroxyalkanoic acid synthase | I | PhaC_N | [1] |
| *Zoogloea ramigera* | P75003 | PHB polymerase | I | PhaC_N | [1] |
| **Species** | **UniProt ID** | **PHA Synthase** | **PhaC Class** | **Pfam Domain** | **Ref** |
| *Pseudomonas aeruginosa* | Q51513 | Class II poly(R)-hydroxyalkanoic acid synthase | II | PhaC_N | [1] |
| *Pseudomonas aeruginosa* UCBPP-PA14 | A0A0H2ZIE8 | Poly(3-hydroxyalkanoic acid) synthase 2 | II | PhaC_N | [2] |
| *Pseudomonas aeruginosa* UCBPP-PA14 | A0A0H2ZIG5 | Poly(3-hydroxyalkanoic acid) synthase 1 | II | PhaC_N | [2] |
| *Pseudomonas chlororaphis* sp. aureofaciens | Q8VV56 | PHA synthase 2 | II | PhaC_N | [1] |
| *Pseudomonas mendocina* | Q8RQ67 | PHA synthase 1 | II | PhaC_N | [1] |
| *Pseudomonas nitroreducens* | Q9AGB4 | PHA synthase 2 | II | PhaC_N | [1] |
| *Pseudomonas oleovorans* | P26494 | Poly(3-hydroxyalkanoate) polymerase 1, PHA polymerase 1 | II | PhaC_N | [3] |
| *Pseudomonas oleovorans* | P26496 | Poly(3-hydroxyalkanoate) polymerase 1 | II | PhaC_N | [1] |
| *Pseudomonas putida* | Q9R9W2 | Class II poly(R)-hydroxyalkanoic acid synthase | II | PhaC_N | [1] |
| *Pseudomonas putida* | Q9R9W4 | Class II poly(R)-hydroxyalkanoic acid synthase | II | PhaC_N | [1] |
| *Pseudomonas putida* DSM 6125 | Q88D23 | Poly(3-hydroxyalkanoate) polymerase 2 | II | PhaC_N | [2] |
| *Pseudomonas putida* | Q8KQ21 | PHA synthase 2 | II | PhaC_N | [2] |
| *Pseudomonas putida* | Q8KQ23 | PHA synthase 1 | II | PhaC_N | [2] |
| *Pseudomonas resinovorans* | Q9X5X7 | PHA synthase 1 | II | PhaC_N | [1] |
| *Pseudomonas resinovorans* | Q9X5X9 | PHA synthase 2 | II | PhaC_N | [1] |
| *Pseudomonas* sp. 61-3 | Q9Z3X9 | PHA synthase 2 | II | PhaC_N | [1] |
| *Pseudomonas* sp. 61-3 | Q9Z3Y1 | PHA synthase 1 | II | PhaC_N | [1] |
| *Pseudomonas stutzeri* | Q848R9 | PHA synthase 2 | II | PhaC_N | [1] |
| *Pseudomonas stutzeri* | Q848S0 | PHA synthase 1 | II | PhaC_N | [1] |
| *Rickettsia prowazekii* Rp22 | D5AY96 | Poly(3-hydroxyalkanoate) synthetase | II | PhaC_N | [4] |
| **Species** | **UniProt ID** | **PHA Synthase** | **PhaC Class** | **Pfam Domain** | **Ref** |
| *Allochromatium vinosum* DSM180^*^ | P45370 | Class III poly(R)-hydroxyalkanoic acid synthase subunit PhaC | III | Abhydrolase_1 | [2] |
| *Allochromatium vinosum* (*Chromatium vinosum*) | Q402A9 | Class III poly(R)-hydroxyalkanoic acid synthase subunit PhaC | III | Abhydrolase_1 | [2] |
| *Chromatium okenii* | A0A2S7XPK6 | Class III poly(R)-hydroxyalkanoic acid synthase subunit PhaC | III | Abhydrolase_1 | [4] |
| *Ectothiorhodospira shaposhnikovii* | Q9F5P8 | Class III poly(R)-hydroxyalkanoic acid synthase subunit PhaC | III | Abhydrolase_1 | [3] |
| *Haloarcula hispanica* DSM 4426^^^ | G0HQZ6 | Class III poly(R)-hydroxyalkanoic acid synthase subunit PhaC | III | Abhydrolase_1  HHH_5 | [2] |
| *Haloarcula marismortui* ATCC 43049^^^ | Q5UYM0 | Class III poly(R)-hydroxyalkanoic acid synthase subunit PhaC | III | Abhydrolase_1  HHH_5 | [2] |
| *Haloferax mediterranei* DSM 1411^^^ | I3R7Z6 | Class III poly(R)-hydroxyalkanoic acid synthase subunit PhaC | III | Abhydrolase_1  HHH_5 | [2] |
| *Haloferax mediterranei* DSM 1411^^^ | I3R9Z4 | Class III poly(R)-hydroxyalkanoic acid synthase subunit PhaC | III | Abhydrolase_1  HHH_5 | [2] |
| *Haloferax mediterranei* DSM 1411^^^ | I3RBD8 | Class III poly(R)-hydroxyalkanoic acid synthase subunit PhaC | III | Abhydrolase_1  HHH_5 | [2] |
| *Halogeometricum borinquense* DSM 11551^^^ | E4NU13 | Poly(R)-hydroxyalkanoic acid synthase, class III, PhaC subunit | III | Abhydrolase_1  HHH_5 | [2] |
| *Halorhabdus utahensis* DSM 12940^^^ | C7NUH3 | Poly(R)-hydroxyalkanoic acid synthase, class III, PhaC subunit | III | Abhydrolase_1  HHH_5 | [2] |
| *Magnetococcus* sp. MC-1 | A0L6T8 | Poly(R)-hydroxyalkanoic acid synthase, class III, PhaC subunit | III | Abhydrolase_1 | [4] |
| *Synechococcus* sp. MA19 | Q8RT55 | PhaC | III | Abhydrolase_1 | [3] |
| *Chlorogloeopsis fritschii* | Q8RTL8 | Class III poly(R)-hydroxyalkanoic acid synthase subunit PhaC | III | Abhydrolase_1 | [2] |
| *Synechocystis* sp. PCC 6803^*^ | P73390 | Class III poly(R)-hydroxyalkanoic acid synthase subunit PhaC | III | Abhydrolase_1 | [4] |
| *Thiocystis violacea*^*^ | P45366 | Poly(3-hydroxyalkanoate) polymerase subunit PhaC | III | Abhydrolase_1 | [3] |
| *Vibrio variabilis* | A0A090SSB5 | Polyhydroxyalkanoic acid synthase | III | PhaC_N | [4] |
| *Xanthomonas campestris* pv. campestris | Q8P8T0 | Poly (3-hydroxybutyric acid) synthase | III | Abhydrolase_1 | [4] |
| **Species** | **UniProt ID** | **PHA Synthase** | **PhaC Class** | **Pfam Domain** | **Ref** |
| *Bacillus anthracis* | A0A384LMA2 | Class III poly(R)-hydroxyalkanoic acid synthase subunit PhaC | IV | Abhydrolase_1 | [3] |
| *Bacillus cereus* E33L | Q63E53 | Poly(R)-hydroxyalkanoic acid synthase, class III | IV | Abhydrolase_1 | [4] |
| *Bacillus megaterium* QMB1551 | D5DZA0 | Polyhydroxyalkanoic acid synthase, PhaC subunit | IV | Abhydrolase_1 | [1] |
| *Bacillus megaterium* | Q9ZF92 | Class III poly(R)-hydroxyalkanoic acid synthase subunit PhaC | IV | Abhydrolase_1 | [2] |
| *Bacillus* sp. INT005 | Q8GI81 | PHA synthase | IV | Abhydrolase_1 | [1] |
| **Species** | **UniProt ID** | **PHA Synthase** | **PhaE** | **Pfam Domain** | **Ref** |
| *Allochromatium vinosum* DSM 180 | P45372 | Poly(3-hydroxyalkanoate) polymerase subunit PhaE | III | PHA_synth_III_E | [3] |
| *Desulfococcus multivorans* DSM 2059 | S7VKE6 | Poly(R)-hydroxyalkanoic acid synthase class III PhaE subunit | III | PHA_synth_III_E | [4] |
| *Ectothiorhodospira shaposhnikovii* | Q9F5P9 | PHA synthase subunit PhaE | III | PHA_synth_III_E | [3] |
| *Haloarcula hispanica* DSM 4426 | G0HQZ5 | Poly(3-hydroxyalkanoate) polymerase subunit PhaE | III | PHA_synth_III_E | [3] |
| *Halogeometricum borinquense* DSM 11551 | E4NU12 | Poly(R)-hydroxyalkanoic acid synthase subunit | III | PHA_synth_III_E | [2] |
| *Magnetococcus marinus* sp. MC-1 | A0L6T7 | Poly(R)-hydroxyalkanoic acid synthase, class III, PhaE subunit | III | PHA_synth_III_E | [4] |
| *Synechocystis* sp. PCC6803 | P73389 | Poly(3-hydroxyalkanoate) polymerase subunit PhaE | III | PHA_synth_III_E | [4] |
| *Thiocystis violacea* | P45367 | Poly(3-hydroxyalkanoate) polymerase subunit PhaE | III | PHA_synth_III_E | [3] |
| *Xanthomonas campestris* DSM 3586 | Q8P8S9 | PHA synthase subunit | III | PHA_synth_III_E | 4 |
| **Species** | **UniProt ID** | **PHA Synthase** | **PhaR** | **Pfam Domain** | **Ref** |
| *Bacillus anthracis* | A0A384L2C0 | PhaR protein | IV | - | [3] |
| *Bacillus cereus* E33L | Q63E55 | PHA synthase subunit PhaR regulator of PhaP | IV | - | [4] |
| *Bacillus megaterium* | Q9ZF94 | PhaR | IV | - | [3] |
| *Bacillus megaterium* QMB1551 | D5DZ98 | Polyhydroxyalkanoic acid synthase, PhaR subunit | IV | - | [1] |
| *Bacillus megaterium* | D2Z0C0 | PHA Synthase | IV | - | [4] |
| *Bacillus* sp. INT005 | Q8GI83 | PHA synthase subunit PhaR | IV | - | [1] |
| *Bradyrhizobium japonicum*^#^ | A0A1L3F0U4 | Polyhydroxyalkanoate synthesis repressor PhaR | IV | PHB_acc_N  PHB_acc | [4] |
| *Bradyrhizobium japonicum* SEMIA 5079^#^ | A0A023XUR3 | Polyhydroxyalkanoate synthesis repressor PhaR | IV | PHB_acc_N  PHB_acc | [4] |
| *Chromatium okenii*^#^ | A0A2S7XQ05 | Polyhydroxyalkanoate synthesis repressor PhaR | IV | PHB_acc_N  PHB_acc | [4] |
| *Chromohalobacter salexigens* DSM 3043^#^ | Q1QZ43 | Polyhydroxyalkanoate synthesis repressor PhaR | IV | PHB_acc_N  PHB_acc | [2] |
| *Magnetococcus marinus* sp. MC-1^#^ | A0L6T5 | Polyhydroxyalkanoate synthesis repressor, PhaR | IV | PHB_acc_N  PHB_acc | [4] |
| **Species** | **UniProt ID** | **PHA Synthase** | **PhaC**  **Class** | **Pfam Domain** | **Ref** |
| *Ralstonia eutropha* H16 | Q0KA68 | Poly(3-hydroxybutyrate) polymerase | PHB Synthase^&^ | PHBC_N  PhaC_N | [2] |
| *Rhodobacter sphaeroides* | Q3IYE8 | Poly(3-hydroxybutyrate) polymerase | PHB Synthase^&^ | PHBC_N  PhaC_N | [4] |

^*^PhaCs in three bacterial species, *Allochromatium vinosum* DSM 180, *Synechocystis* sp. PCC 6803, and *Thiocystis violacea* are classified as Class I PHA synthase while three other papers classify these enzymes as Class PHA synthases, which is more accurate due to their comparatively short sequence lengths. In fact, names and classification of PhaCs in the literature are still ambiguous and require further investigation.

^^^A special group of Class III PHA synthase due to the presence of HHH-5 domain, which is exclusively associated PhaCs in halophilic organisms.

^#^A special group of PhaR due to the presence of PHB_acc_N and PHB_acc domains. Normally, PhaR does not have a specified domain.

^&^A special group of PhaCs for exclusive PHB synthesis.
